# Expanding the Evo-Devo Toolkit: Generation of 3D mammary tissue from diverse mammals

**DOI:** 10.1101/2023.06.28.546911

**Authors:** Hahyung Y. Kim, Ishani Sinha, Karen E. Sears, Charlotte Kuperwasser, Gat Rauner

**Author notes:** Corresponding Authors: Gat Rauner, Tufts University School of Medicine, Department of Developmental, Chemical & Molecular Biology, 136 Harrison Ave., Boston, MA 02111. Charlotte Kuperwasser, Tufts University School of Medicine, Department of Developmental, Chemical & Molecular Biology, 136 Harrison Ave., Boston, MA 02111.

## Abstract

The divergent events during mammary gland development between species and across evolution are not well studied mainly due to the lack of tractable model systems. In theory, advancements in the field of organoid technology now make it possible to study developmental processes adapted throughout species evolutions to accommodate advantageous phenotypes. However, its application to the mammary gland remains limited to rodents and humans.

In the current study, we successfully created next-generation 3D mammary gland organoids from eight eutherian mammals and the first 3D organoid of a marsupial mammary gland (the gray short-tailed opossum), representing a more ancient version of the mammary gland. Using mammary organoids, we identified a role for ROCK protein in regulating branching morphogenesis, a role that manifests differently in human and opossum mammary organoids. This finding demonstrates the utility of 3D organoid model for understanding the evolution and adaptations of signaling pathways.

Overall, the establishment of mammary organoids as animal model surrogates is a significant advancement in the field of mammary gland biology and evolution and paves the way for future studies utilizing this model. This achievement highlights the potential of organoid models to expand our understanding of mammary gland biology and evolution and their utility in studying milk production and breast cancer research.

## INTRODUCTION

The emergence of mammals in evolution is marked by the appearance of a mammary gland: a secretory gland able to provide nutrition to offspring. The mammary gland evolved from a cutaneous, likely hair-associated gland, with evidence pointing to the apocrine sweat gland as the precursor of the mammary gland [1], and has undergone remarkable evolutionary transformations, adapting to diverse ecological pressures and reproductive strategies.

The earliest extant mammals are the monotremes, consisting of only two: the echidnas and the platypus. Monotremes are egg-laying mammals that diverged from the mammalian class about 200 million years ago [2] (Figure 1A). The monotreme mammary gland is a patch of special hairs, where milk is secreted from a gland at the base of each hair, of which there are approximately 100-150 in the echidna [3].

**Figure 1.**
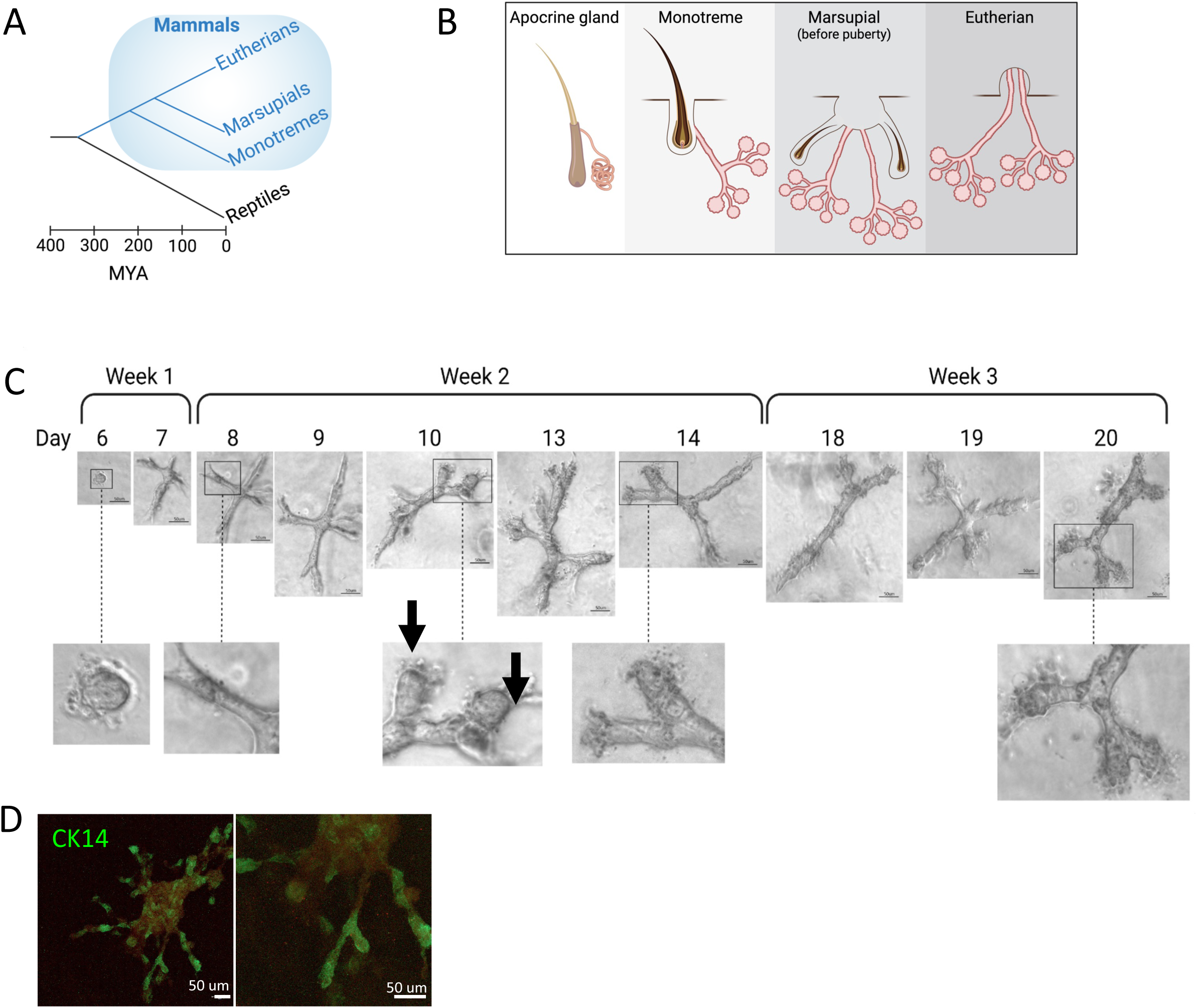
Generation of marsupial mammary gland organoids. **A.** The timeline of mammalian group divergence in evolution. **B.** Illustration of mammary gland structure in each of the three mammalian groups, compared with an apocrine gland. **C.** Bright field images depicting the development of opossum mammary gland organoids during 3 weeks in culture. **D.** Whole mount in-situ hybridization detection of keratin 14 in 3D opossum organoids.

The next mammalian group to diverge is the marsupials. Marsupials, also known as Metatheria, diverged from placental mammals 160 million years ago, or more [2, 4, 5]. The marsupial mammary gland develops associated with a hair follicle (that disappears in the adult gland), representing an ancient version of a gland that evolved from the cutaneous apocrine gland [1]. Marsupial young are born altricial, relying less on placentation and completing their development postnatally, with an extended period of lactation. Marsupials comprise about 10% of mammals, and the rest are eutherian mammals (also known as “placental mammals”).

The mammary gland of eutherians does not develop in association with hair and includes a nipple. Eutherian mammals are the most taxonomically diverse group of mammals, with over 4000 species representing a large variety of mammary gland morphologies, milk compositions, and lactation strategies, that evolved throughout mammalian evolution (Figure 1B).

Investigating the evolution of the mammary gland can offer invaluable insights into fundamental biological processes, including tissue development, hormonal regulation, lactation, and immunity. The exploration of mammary gland evolution not only contributes to comparative anatomy and phylogenetics, unraveling the evolutionary relationships among mammalian species, but also has profound implications for biomedical research. Certain species have evolved unique lactation adaptations, which can serve as valuable models for understanding the regulation of milk composition and the cyclical regeneration of mammary glands. Understanding the genetic and molecular mechanisms that enable these adaptations can provide crucial knowledge for advancements in breast cancer research, reproductive medicine, and the development of therapies for lactation-related disorders.

One of the primary obstacles to deep understanding of mammary gland evolution is the limited access to live specimens, the ethical concerns associated with conducting invasive experiments on animals, and the unsuitability of most species as laboratory animals. Traditional approaches relied heavily on comparative anatomy and histological analysis of preserved tissues, which provided valuable but static information. However, these methods often fell short in capturing the dynamic nature of mammary gland development, lactation, and tissue regeneration. Furthermore, the functional and molecular aspects of mammary gland biology, including the regulation of milk composition and the intricate cellular processes underlying cyclical regeneration, remained elusive.

Recent advances in organoid technology enabled the generation of 3D mammary gland tissue from single primary cells isolated from an adult human breast [6, 7]. The orchestrated process of development from single cells to a complex network of ducts and lobules comprised of multiple epithelial cell types recapitulates important aspects of mammary tissue development. It, therefore, provides a valuable model to gain a deep understanding of the events and conditions that drive mammary gland development and regeneration, at the cellular and molecular levels. Importantly, it enables the deciphering of developmental programs that guide stem and progenitor cells in making cell fate decisions.

Thus far, complex ductal-lobular 3D organoids of the mammary gland, that capture both the branched and acinar components of the milk duct network, were generated from adult mammary cells of humans and mice and contributed to our understanding of mammary gland development and regeneration in these species [8, 9]. However, 3D organoid models of the mammary gland capturing these morphologies were not previously generated from other species. In this work, we endeavored to grow mammary gland organoids from 8 eutherian mammals, and one marsupial. We show that individual cells isolated from adult mammary glands of these species proceed to form branched epithelial structures in a 3D matrix. This report includes the first 3D organoid of a marsupial mammary gland: the gray short-tailed opossum.

Furthermore, we identify that the ROCK inhibitor Y-26732 is necessary for the branching of opossum, but not human, mammary organoids in a 3D collagen matrix. Interestingly, human breast organoids exhibited a hyperbranched phenotype when treated with ROCK inhibitor. This finding indicates that ROCK plays a role in regulating branching morphogenesis, that this role may be conserved in early mammals, but that it may manifest differently in different mammalian groups or species.

Importantly, this report describes the establishment of 3D mammary organoids as surrogates of animal models previously unavailable for mammary gland biology research. This organoid model can serve studies focused on evolutionary and developmental biology, as well as milk production and breast cancer research.

Ultimately, the study of mammary gland evolution not only illuminates the biological history of mammals but also unveils the physiological mechanisms that underlie crucial aspects of mammalian life, offering significant insights for improving human health.

## RESULTS

### 3D marsupial mammary gland organoids from the gray short-tailed opossum

The gray short-tailed opossum, *monodelphis domestica*, is a pouch-less marsupial that is long studied as an animal model of early mammals [10, 11]. The gray short-tailed opossum mammary gland has 13 nipples: 12 arranged in a circle around one [12-14]. The adult marsupial mammary gland includes an intra-mammary muscle, the ilio marsupialis, unique to marsupials, that can be seen near the mammary epithelium [15] (Supplemental figure 1A). Surrounding the epithelium is an area of fibrotic tissue, as seen in some eutherian mammals.

We aimed to grow mammary gland organoids of the opossum mammary gland, which could serve as a model to study mammary gland development in a marsupial mammal. Primary mammary epithelial cells were isolated from an adult (17 months old), virgin, female gray short-tailed opossum, and embedded in 3D gels of either Matrigel or ECM hydrogel, supplemented with MEGM, as previously described for the generation of human and mouse mammary organoids [6, 16, 17].

The organoids that developed were acinar and failed to branch after 13 days in culture (Supplemental Figure 1B), indicating that the conditions sufficient for the growth of human breast organoids did not support the full maturation of the opossum mammary epithelium. To try to induce organoid branching morphogenesis, we changed the matrix composition to include Matrigel in addition to collagen. Matrigel is rich in laminins and can successfully support the growth of mammary organoids, as well as organoids of other tissues. We included Matrigel at a range of percentages (0, 25, 50, 75, and 100) in the hydrogels while maintaining the balanced ratio of the other components of the gel, as shown in Supplemental Figure 1C. Branching was not observed in any of the gel compositions (Supplemental Figure 1D). We observed that collagen gel combined with Matrigel has altered consistency and is opaquer than collagen alone or Matrigel alone. This could result from interference with the collagen’s polymerization process, causing changes in the physical properties of the gel. To avoid this, we increased laminin concentration in the hydrogel by adding laminin directly, without Matrigel. Increasing the laminin concentration 5- or 10-fold (to final concentrations of 125 ug/mL and 250 ug/mL, respectively), did not result in the branching of opossum mammary organoids (Supplemental Figure 1E).

We attempted to induce organoid branching by adding factors to the media that may induce branching. Estrogen and progesterone regulate the expansion of the mammary epithelium, and we added them to the media alone or in combination. Prostaglandin E2 induces aromatase, which is necessary for estrogen biosynthesis [18], and its receptors are expressed during pregnancy and lactation in the mouse mammary gland, indicating a role in tissue development [19]. Adding either of those to the media did not lead to the branching of the opossum mammary epithelial organoids (Supplemental Figure 1F). Taken together, the failure of our multiple attempts to address the lack of ductal elongation and maturation of opossum organoids indicates that it requires significantly different growth conditions compared with human or mouse mammary organoids and may reflect differences in the developmental biology of this more primitive mammary gland.

We then attempted to culture opossum mammary epithelial cells in commercial organoid media (Intesticult, Cat# 06010, StemCell Technologies, Vancouver, Canada). The formulation of the media is proprietary but is based on published studies that optimized it for intestinal epithelium organoids [20, 21]. Media supplements described in these studies have been reportedly used in breast organoid culture medium [22, 23], Table 1). Remarkably, opossum organoids developed and branched in Intesticult (Figure 1C). Branching was visible after 7 days, and enlargement of the tips of branches resembling lobules was visible by day 10. However, not all branches had such lobular enlargements even after 3 weeks in culture, and mature lobules such as seen in the culture of human breast organoids [6] were not seen during this time. In-situ hybridization for opossum CK14 demonstrated expression in the basal layer of the organoid, as seen in eutherian species [24-26] (Figure 1D).

**Table 1.** Media supplements for intestinal epithelial organoids that are also used in breast organoid media formulations.

| **Intestinal organoid media supplement** | **Used in breast organoid media** |
| --- | --- |
| [Leu<sup>15</sup>] Gastrin-I | - |
| A-83-01 | + |
| B27 Supplement | + |
| DNase | - |
| EDTA | - |
| GlutaMAX-I | + |
| Human recombinant FGF10 | + |
| Human recombinant R-spondin | + |
| Mouse recombinant EGF | + |
| Mouse recombinant Noggin | + |
| Mouse recombinant Wnt-3A | + |
| N2 Supplement | - |
| N-acetylcystein | + |
| Nicotinamide | + |
| SB202190 | + |
| Y-27632 | + |

### ROCK inhibition is required for the branching of opossum mammary organoids

Next, we attempted to identify which factors in Intesticult were necessary for the formation of opossum organoids, by selectively adding to MEGM, individually or in groups, supplements used in the studies that the Intesticult formulation is based on [20, 21]. Figure 2A lists the ingredients included in each of the combinations we tested, and the representative result of each. In all experiments, we used MEGM without supplements and the commercial Intesticult media as negative and positive controls. We identified the commercial supplements B27 and N2 as necessary for the formation of opossum mammary gland organoids. We also identified that a group of 3 inhibitors were necessary for growth to occur: Y-27632, an inhibitor of Rho-associated, coiled-coil containing protein kinase (ROCK) [27]; SB202190 a p38 MAPK inhibitor [28]; and A83-01, an inhibitor of activin receptor-like kinases ALK5, ALK4, and ALK7 [29].

**Figure 2.**
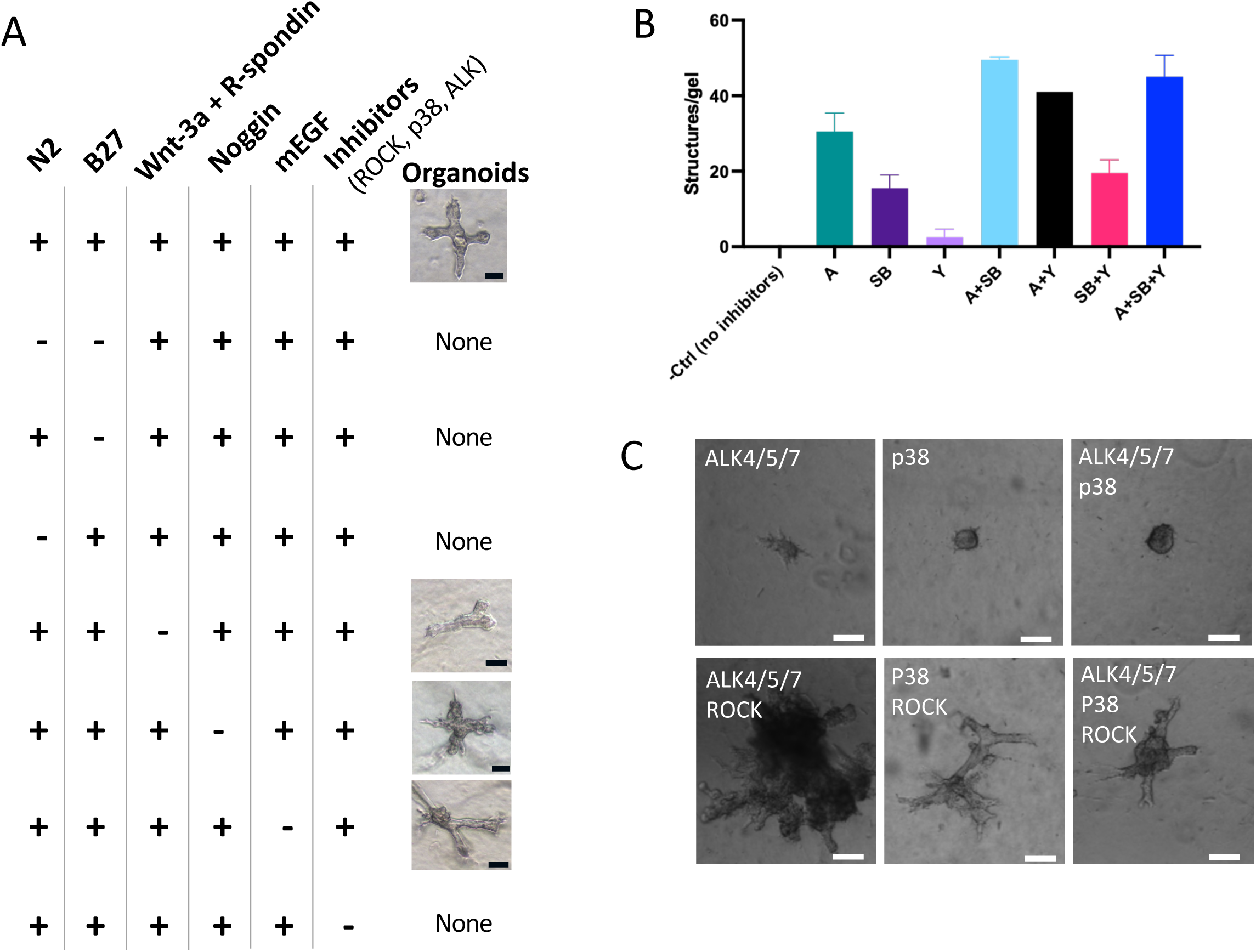
Dissecting the minimal media for marsupial organoid development. **A.** Table showing the combinations of media supplements and representative bright field images of opossum organoids that developed in each. Scale bar = 50 μm. **B.** Quantification of the number of organoids that developed per gel in the presence of combinations of specific chemical inhibitors. “A” is A83-01; “SB” is SB202190; “Y” is Y-27632. Error bars represent STDEV. **C.** Representative bright field images of opossum organoids that developed in the presence of specific chemical inhibitors. Scale bar = 50 μm.

To find which of the inhibitors was necessary for growth, we added them to the media separately or in combination and quantified the number of structures formed by each of the combinations (Figure 2B). We concluded that A83-01 and SB202190 were each necessary for organoids to form and that together they induced a synergistic effect of increasing the number of organoids. The ROCK inhibitor Y-27632 was not necessary or sufficient for organoid formation, but observing organoid morphology indicated that branching only occurred when Y-27632 was present (Figure 2C). Based on these experiments, we formulated a defined media for the culture of opossum organoids, which includes the supplements required for growth, and Y-27632 required for branching (Table 2).

**Table 2.** Opossum mammary organoids media

| Reagent | Media concentration | Supplier | Catalog number |
| --- | --- | --- | --- |
| B27 | 1x | Thermo Fisher Scientific | 17504044 |
| N2 | 1x | Thermo Fisher Scientific | 17502048 |
| SB202190 | 10 uM | Thermo Fisher Scientific | J60950.MB |
| Y-27632 | 10 uM | Bio-Techne | 1254 |
| A-83-01 | 500 nM | AG Scientific | A-1764 |
| Thermo Fisher Scientific, Waltham, MA; Bio-Techne, Minneapolis, MN; AG Scientific, San Diego, CA |  |  |  |

Our previous experimental observations indicated that human breast organoids do not require the addition of ROCK inhibitors to facilitate their branching [6, 7]. ROCK inhibitors are routinely added to human primary cell cultures in many protocols, including some protocols for the culture of primary mammary epithelial cells, to mitigate cell death due to anoikis and maximize cell viability [30, 31]. We cultured human and opossum mammary organoids under the same matrix and media conditions, to see if they exhibit a different response to the presence of ROCK inhibitor. As expected, opossum organoids branched only in the presence of a ROCK inhibitor, while human organoids branched in its absence (Figure 3A, B). Interestingly, the presence of a ROCK inhibitor resulted in the hyper branching of human organoids. The hyperbranched phenotype we observed in the human organoid is consistent with the role of ROCK in facilitating branching morphogenesis by maintaining polarization within bi-layered epithelial tissues, as reported by others [32, 33].

**Figure 3.**
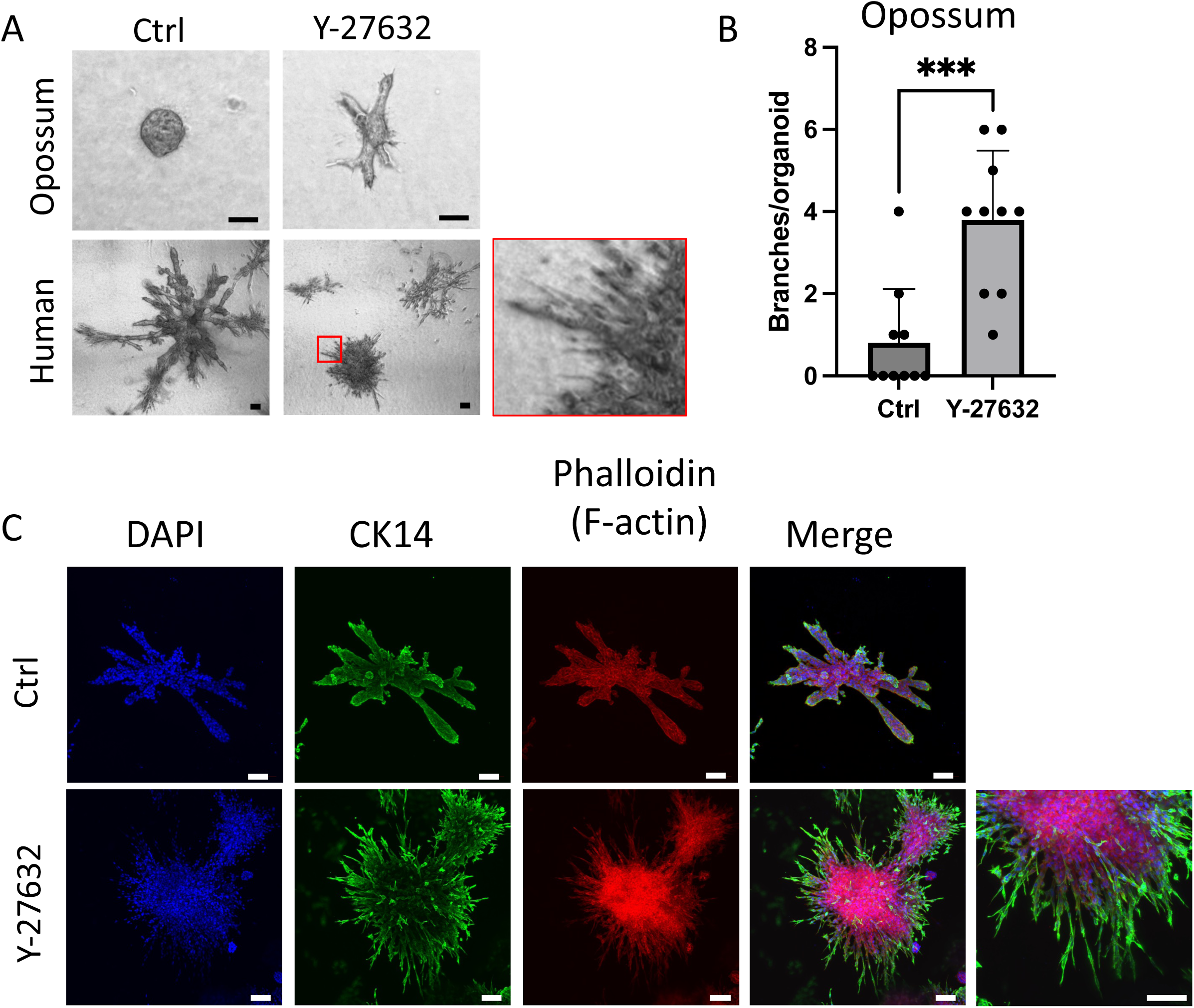
ROCK inhibition is necessary for the branching of opossum mammary organoids but disrupts the branching of human counterparts. **A.** Bright field images of representative human and opossum mammary gland organoids that developed with or without the ROCK inhibitor Y-27632. Scale bar = 50 μm. **B.** Quantification of branches per organoid in opossum mammary organoids with or without Y-27632. Error bars represent STDEV. **C.** Immuno-fluorescent staining of human breast organoid with or without Y-27632 for CK14 (green) and F-actin (phalloidin, red). Scale bar = 100 μm.

We stained human breast organoids cultured with or without Y-27632 for F-actin and the basal marker CK14 (Figure 3C). Our findings revealed that the administration of a ROCK inhibitor led to the emergence of CK14-positive cells extending outward from a densely clustered core of epithelial cells in the organoid. This starkly contrasted with the presence of a single layer of CK14-positive cells enveloping a branched epithelial structure in the untreated organoids.

### Generation of 3D mammary gland organoids from 8 Eutherian mammals

Mammary gland tissue samples were collected from 8 eutherian species, representing four mammalian orders: Carnivora (ferret and cat), Artiodactyla (pig, goat, and cow), Lagomorpha (rabbit), and Rodentia (rat and hamster) (Figure 4A). Single primary cells from each species were embedded in ECM hydrogels, as previously described for the generation of human breast organoids [6]. Organoids from all species were cultured for 11-13 days in identical organoid media (Intesticult).

**Figure 4.**
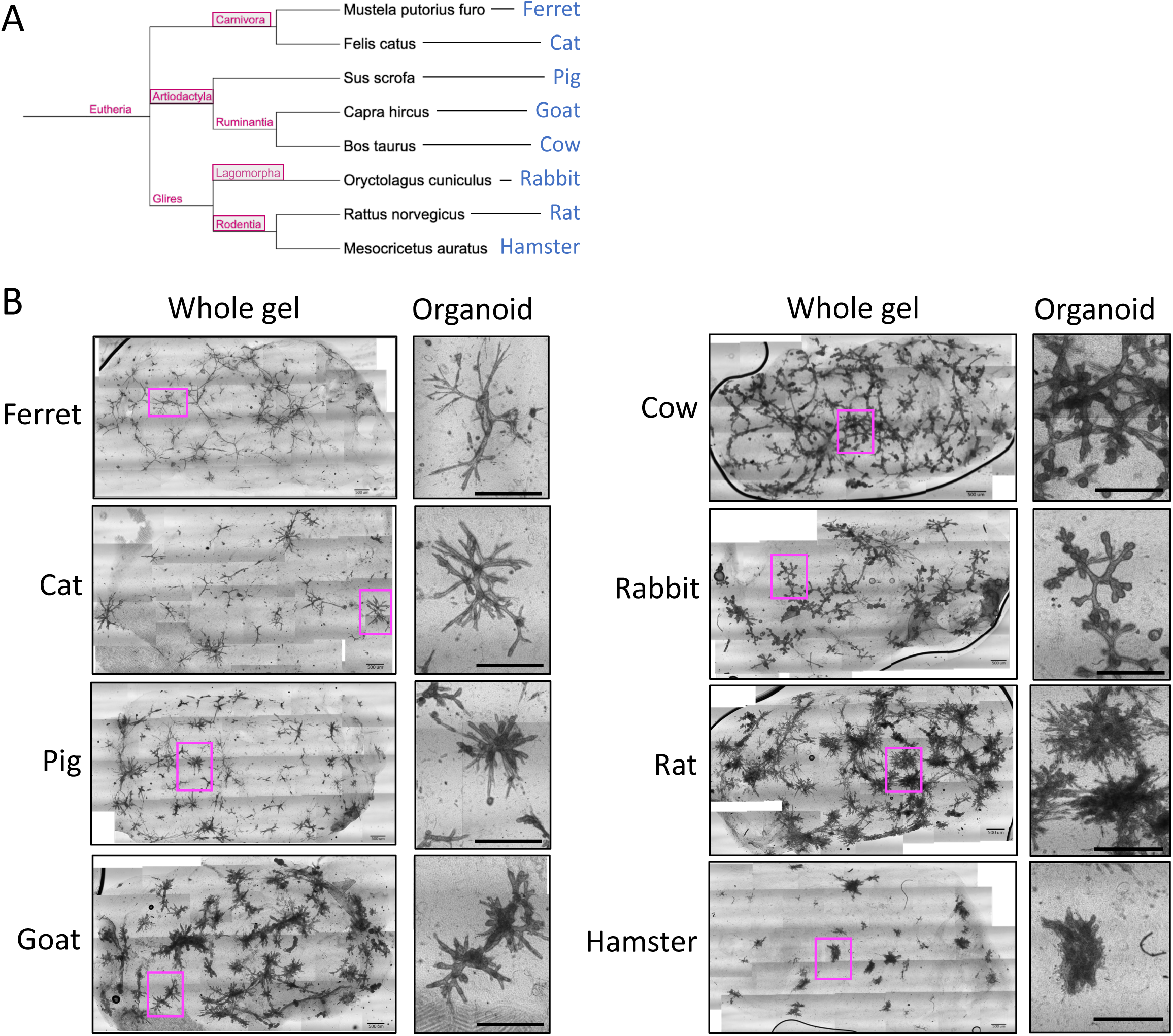
Generation of mammary gland organoids from diverse eutherian mammals. **A.** Phylogenetic tree depicting the common ancestry of the 8 eutherian mammals that participated in the study. **B.** Brightfield images of hydrogels occupied by mammary tissue organoids from 8 eutherian species. Scale bar = 0.5 mm.

3D organoids were successfully generated from primary mammary epithelial cells from all 8 eutherian mammals (Figure 4B). Varied morphological characteristics were observed between species. Specifically, while branching morphogenesis occurred in organoids from all species analyzed, distinctly shorter ductal elongation was observed for the hamster. Defined lobules were visible in organoids of cow, goat, pig, and rabbit (artiodactyls and lagomorphs), but not in organoids of cat, ferret, hamster, and rat (carnivores and rodents).

We stained the organoids with phalloidin, revealing the distribution of filamentous actin (F-actin) in the epithelial structures that formed (Figure 5A). F-actin encapsulated the epithelial structures in organoids from cow, rabbit, cat, and pig, but not in organoids from ferret, rat, goat, and hamster, in which phalloidin stained the organoid mostly internally.

**Figure 5.**
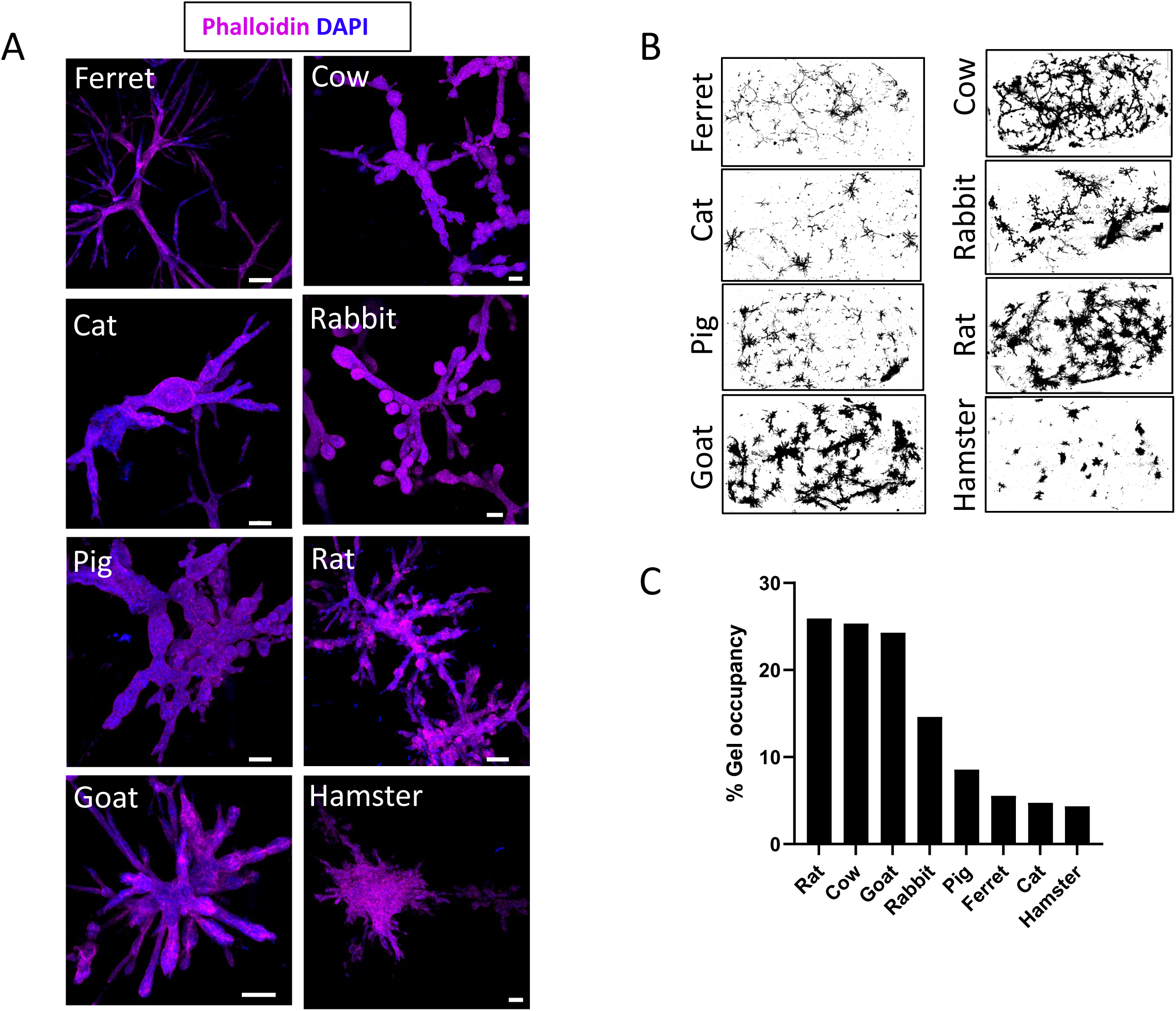
Comparative analysis of mammary organoids from eutherian mammals. **A.** Fluorescent staining for DAPI (nuclei, blue) and phalloidin (F-actin, magenta) of representative organoids from each species. Scale bar = 100 μm **B.** Thresholded images of gels occupied by organoids used to measure gel occupancy. **C.** Graph depicting gel occupancy by organoids from each of the species.

We observed the convergence of adjacent organoids, forming a network in the gel, particularly in cow, rat, and rabbit organoids. There was also a difference in the degree to which the organoids filled the gel matrix, impacted by differences in the thickness of the organoids and their ducts and lobules, as well as the number of organoids that developed in the gel (Figure 5B, C). The percentage of gel occupancy was highest in gels seeded with rat, cow, and goat cells (25.9%, 25.3%, and 24.3%, respectively), and lowest for the ferret, cat, and hamster (5.5%, 4.7%, and 4.3%, respectively). Rabbit and pig organoids gel occupancy fell in between, with 14.6% and 8.6%, respectively.

Overall, the results demonstrate that mammary gland organoids can be generated from a wide range of eutherian mammals. These organoids can provide a valuable model for the study of mammary gland development and disease in a variety of species.

## DISCUSSION

In this work, we generate 3D mammary gland tissue organoids from multiple eutherian mammals and one marsupial. We showed that individual cells isolated from adult mammary glands of these species proceeded to form branched epithelial structures in a 3D matrix. These organoids can serve to gain a deeper understanding of the events and conditions that drive mammary gland development and regeneration, at the cellular and molecular levels, across mammals.

The successful generation of mammary organoids from these species using the same culture conditions suggests that mammary gland development and branching morphogenesis are regulated by conserved molecular mechanisms across mammalian species, while at the same time, the role of certain signaling pathways may differ between species. Particularly, the study identified that ROCK inhibition is required to enable branching morphogenesis of opossum mammary organoids, while their human counterparts branched in the absence of such inhibition, and branching was perturbed if ROCK was inhibited. This implies a differential role for ROCK in branching morphogenesis in different species. Further studies are required to determine what mechanisms underlie the difference, and whether it represents a difference between an evolutionarily ancient vs. a more recent version of the gland, or is manifested differently in different species regardless of their evolutionary age.

A crucial aim in evolutionary developmental biology is to comprehend the emergence and progression of the molecular mechanisms that support the development of tissues and organisms. The Rho/ROCK pathway controls actin cytoskeletal organization and plays a role in cell movement and shape in single-cell organisms such as the amoeba [34], in the development of sponges [35], and in complex cellular functions that are important for development, such as cell differentiation and morphogenesis, in vertebrates [36-38]. A role for ROCK proteins in the branching morphogenesis of epithelial tissues, specifically, was identified in the mouse mammary gland and submandibular gland (SMG), and in rat kidneys [32, 33, 39]. In these studies, ROCK signaling mediated branching morphogenesis via controlling cell polarization in the bi-layered epithelia. Inhibition of ROCK resulted in reduced budding of the SMG during embryonic development, led to the formation of poorly patterned branched organoids of mammary epithelium, and to altered branching morphogenesis in the rat kidney. The results of these studies indicate that ROCK signaling is required for the appropriate branching of bi-layered epithelia in the mouse. This is in line with our findings of hyperbranched human mammary organoids upon ROCK inhibition, reflecting dysregulation of the branching process. The versatility of the ROCK pathway has the potential to be adaptable across species, and it can be hypothesized that it mediates branching differently, depending on the context. Our findings indicate that ROCK inhibition enables branching of opossum mammary organoids, under the matrix and media conditions we tested. While this could reflect a differential role of ROCK between species (human/opossum) or between mammalian groups (eutherian/marsupial), it could also reflect differences in the response of cells from those species to the matrix conditions. Indeed, it was shown that the ROCK pathway impacted branching orientation in mammary organoids cultured in a randomly oriented collagen matrix, but not in a positionally aligned matrix [40].

It would be valuable to expand the generation of mammary epithelial organoids to include more species to gain a more comprehensive understanding of mammary gland development across mammals. It would be particularly valuable to include additional marsupials in the analysis, to determine if this group of mammals exhibits unique regenerative or other mechanisms of mammary epithelium.

Of particular interest are species that rarely develop cancer of the mammary gland (e.g., cows, pigs, goats, and others [41-45]), or develop aggressive mammary tumors (e.g., cats [46]). Agricultural research into the regulation of milk production and milk composition in dairy animals such as cows and goats would also benefit greatly from a 3D ex-vivo model.

Importantly, mammary organoids from species not included here, including those with unique lactation adaptations and lactation strategies [47-49], may also be cultured similarly in 3D and provide insight into the molecular and cellular mechanisms that support these adaptations. Finally, a 3D organoid model of the mammary gland from a variety of species can enhance accessibility to animal models previously unavailable for studies of mammary gland biology, from basic and developmental research to milk production and cancer research.

The use of 3D mammary organoids as surrogates for animal models previously unavailable for mammary gland biology research opens new avenues for studying the evolution and development of mammary glands, and lactation biology, as well as investigating the mechanisms of breast cancer development and potential treatments. In the context of cancer, the long-recognized phenomenon of natural resistance to mammary cancer, observed in some species like cows, pigs, and horses, but also ferrets and others, can now be explored using organoids. Despite being known for decades, research into the mechanisms underlying this phenomenon is extremely limited, not least because these large or wild animals are challenging to study in a laboratory setting. The availability of mammary organoids from species that are resistant and susceptible to mammary cancer can enable the study of natural mechanisms of resistance to tumorigenesis of the mammary gland. The agricultural and dairy industries could benefit from the development of mammary gland organoids from different mammalian species, as they could provide insights into milk composition and lactation strategies across different species. This knowledge could inform breeding and management practices to improve milk production and quality, as well as animal welfare.

While the study’s findings contribute significantly to the understanding of mammary gland development and regeneration, there are some limitations that should be acknowledged and addressed in future work. Importantly, the composition of the matrix and media for each species can be modified to better reflect the in-vivo conditions of the tissue, as was successfully done previously for ECM of human breast organoids [6], and here for the medium of opossum organoids. This line of research is likely to identify optimal ECM conditions that support the maturation of tubular-ductal mammary gland tissue in different species, thereby identifying differential stromal-epithelial crosstalk pathways among mammals. In addition, samples from each species, representing variation in factors such as age and parity would increase the value of organoids as surrogates of the original tissue.

To conclude, this paper describes a key advancement in the field of mammary gland biology, namely the successful generation of 3D mammary gland tissue organoids from multiple mammalian species, including one marsupial. This advancement allows for the investigation of mammary gland development, regeneration, and lactation biology across different species, which can lead to a deeper understanding of the conserved molecular mechanisms involved in mammary gland development in mammals, as well as the identification of unique regenerative or other mechanisms of mammary epithelium in certain species.

## MATERIALS AND METHODS

### Sample Collection and Processing

Mammary gland tissue from eight eutherian mammals (rabbit, rat, dog, pig, goat, ferret, hamster, and cow) was obtained under aseptic conditions, from non-lactating, clinically healthy females of each species (BioChemed Services, Winchester, VA, USA). Mammary gland tissue of gray short-tailed opossum was collected from a 17-month-old virgin female, from the colony of the Sears lab at UCLA. Human breast tissue that would otherwise have been discarded as medical waste following reduction mammoplasty surgery was obtained in compliance with all relevant laws, using protocols approved by the institutional review board at Tufts Medical Center. All tissues were anonymized before transfer and could not be traced to specific patients; for this reason, this research was provided exemption status by the Committee on the Use of Humans as Experimental Subjects at Tufts University Health Sciences (IRB# 13521). All patients enrolled in this study signed an informed consent form to agree to participate in this study and for publication of the results.

Tissues were delivered either promptly upon collection or shipped overnight in a saline buffer on ice and immediately processed in the lab. Tissue processing followed the protocol previously described for human breast tissue [7]. If present, surrounding adipose tissue was removed and discarded and the glandular tissue was minced with scissors into ∼1 mm^3^ pieces. Tissue was weighed and incubated in an enzyme solution containing 1.5 mg/mL collagenase A (Sigma Aldrich, Cat# 11088793001) and 0.3 mg/mL hyaluronidase (Sigma Aldrich Aldrich, St. Louis, MO, USA, Cat# H3506) in MEGM (Lonza, Basel, Switzerland, Cat# CC-3150), at a ratio of 10 mL enzyme solution per up to 2 g tissue. Tissues were then incubated rotating for 3 hours at 37°c. Following incubation, tubes were rested upright for 5 minutes to allow epithelial tissue fragments to sink to the bottom. Fat that accumulated on the top was removed using a pipette, and the intermediate liquid containing the stromal fraction (SF) was collected separately from the epithelial fragments at the bottom of the tube. The SF and epithelial fragments were washed 3 times in PBS to remove the enzyme solution, resuspended in MEGM supplemented with 10% DMSO (Fisher Scientific, Waltham, MA, USA, Cat# BP231-100), and cryopreserved in aliquots of 1 mL.

### Organoid Generation and Culture

Cryopreserved samples of enzymatically digested tissue were thawed and further dissociated with trypsin and dispase to obtain a single-cell suspension of mammary epithelial cells, as previously described [7]. Briefly, after washing in MEGM to remove DMSO, the samples were resuspended in pre-warmed 0.25% trypsin-EDTA (Thermo Fisher Scientific, Waltham, MA, USA Cat# 25200-056) and incubated at 37°c for 5 minutes. A defined trypsin inhibitor (Thermo Fisher Scientific, Cat# R007100) was added at 2x trypsin volume. The Sample was centrifuged for 4 minutes at 300 x g, and the pellet was resuspended in 5mg/mL dispase II solution (Sigma Aldrich, Cat# 04942078001) supplemented with 0.1 mg/mL DNase I (Sigma Aldrich, Cat# 4716728001) to dissolve stringy DNA. Following 2 minutes of incubation in dispase-DNase solution, the suspension was filtered through a 40 μm cell strainer (VWR, Radnor, PA, USA, Cat# 76327-098) to yield a single-cell suspension. Cells were counted with trypan blue (Thermo Fisher Scientific, Cat# 15250-061) using a cell counter (TC-20 from BioRad, Hercules, CA, USA), and a cell suspension was prepared at a concentration of 0.5*10^6^ cells/mL. Hydrogels were prepared as previously described [7]. Collagen type I (rat tail, Sigma Aldrich, Cat# 08-115) was mixed on ice with cold MEGM to a final concentration of 1.7 mg/mL and supplemented with 0.1 N NaOH at a volume that equals 12.5% of the collagen volume. Laminin (Thermo Fisher Scientific, Cat# 23017015), fibronectin (Sigma Aldrich, Cat# F1056), and hyaluronic acid (Sigma Aldrich, Cat# 385908) were added to the collagen mix to yield a final concentration of 25 μg/mL, 25 μg/mL, and 12.5 μg/mL, respectively. Cell suspension in MEGM was added to a final concentration of 2.5*10^4^ cells/mL. Hydrogel solution was mixed and deposited as 200 μL gels in 4-well chamber slides (Corning, Corning, NY, USA, Cat# 354104), and allowed to solidify at 37°c for 1 hr. After the hydrogels solidified, 1 mL media was added per well, and the gels were mechanically detached from the well surface using a pipette tip. Mammary organoids from eutherian species were cultured in Intesticult human organoid growth media (StemCell Technologies, Vancouver, Canada, Cat# 06010). Mammary organoids of opossum were cultured in either Intesticult, or opossum organoid media, as described in Table 2. In the relevant experiments, media supplements were added at the concentrations indicated in Table 3. Media was changed 3 times a week. Primary cultures of opossums are maintained at 32.5°C, as previously described for organotypic cultures derived from developing opossum cortex because they have a lower body temperature than most placental mammals and other marsupials [50-52].

**Table 3.** Media supplements for the growth of opossum mammary organoids

| Reagent | Final concentration | Supplier | Catalog number |
| --- | --- | --- | --- |
| A-83-01 | 500 nM | AG Scientific | A-1764 |
| B27 | 1x | Thermo Fisher Scientific | 17504044 |
| Estrogen | 10 ng/mL | Sigma Aldrich | E2257 |
| Human recombinant R-spondin | 1 ug/mL | Thermo Fisher Scientific | A42586 |
| Mouse recombinant EGF | 50 ng/mL | Thermo Fisher Scientific | PMG8041 |
| Mouse recombinant Noggin | 100 ng/mL | Peprotech, Cranbury, NJ | 250-38 |
| Mouse recombinant Wnt-3A | 100 ng/mL | Peprotech | 315-20 |
| N2 | 1x | Thermo Fisher Scientific | 17502048 |
| Progesterone | 500 ng/mL | Sigma Aldrich | P8783 |
| Prostaglandin E2 | 1μM | Sigma Aldrich | P0409 |
| SB202190 | 10 uM | Thermo Fisher Scientific | J60950.MB |
| Y-27632 | 10 uM | Bio-Techne | 1254 |

### Immunofluorescence Staining

Gels were fixed by 30 minutes rotation incubation at room temperature in freshly prepared 4% paraformaldehyde (Electron Microscopy Sciences, Hatfield, PA, USA, Cat# 15710S) diluted in PBS-T, followed by permeabilization overnight in rotation at room temperature in 0.1% Triton X-100 in PBS-T. DAPI and Phalloidin staining was performed by incubating gels for 30 minutes in 2 μg/mL DAPI (Thermo Fisher Scientific, Cat# D1306) and Phalloidin-iFluor 647 (Abcam, Cambridge, UK, Cat# ab176759) according to manufacturer protocol. Primary antibody for CK14 (Thermo Fisher Scientific, Cat# RB-9020-P1) was diluted 1:300, incubated overnight at 4°C, followed with 2 hour incubation with anti-rabbit Alexa-fluor 488 conjugated secondary antibody (Thermo Fisher Scientific, Cat# A-11008) diluted 1:500.

### Whole mount in-situ hybridization

Whole mount in-situ hybridization (WISH) for the detection of opossum keratin 14 was performed using HCR RNA-FISH (Molecular Instruments, Los Angeles, CA, USA). Fluorescently labeled probes for *monodelphis domestica* KRT14 were planned based on the ensemble transcript ENSMODT00000074233, and the hybridization and staining was performed according to the manufacturer’s protocol. Fixation of hydrogels for WISH was performed in 4% paraformaldehyde in sterile PBS for 1 hour, followed by 3 washes in sterile PBS-T, dehydration in methanol for 5 min. on ice, repeated twice. Hydrogels were stored in methanol at -20C overnight, and rehydrated with a series of methanol in PBS-T (75%, 50%, 25%), each for 5 minutes on ice, followed by two washes with PBS-T. hydrogels were then incubated with proteinase K (10 µg/mL in PBS) for 3 minutes at room temperature, and washed for 5 minutes with PBS-T. Fixation in 4% PFA was repeated, for 20 minutes at room temperature, followed by two washes in PBS-T on ice. Subsequently, hydrogels were washed in 1:1 PBS-T : 5x SSCT buffer for 5 minutes on ice. 5x SSCT buffer was prepared by diluting 20x SSC (saline sodium citrate) in DEPC water and adding 0.1% tween (Sigma Aldrich, Cat# P1379). Next, hydrogels were washed in 5x SSCT buffer for 5 minutes on ice. Hydrogels were then incubated with pre-hybridization and probe solution, washed, and incubated with the amplification reagents, as indicated by the kit protocol.

### Imaging and analysis

Bright-field images were obtained using a Nikon Eclipse Ti-U inverted microscope (Nikon Instruments, Melville, NY, USA) equipped with a digital camera and matching software (SPOT Imaging, Sterling Heights, MI, USA). Full gel images were manually assembled from fields imaged sequentially. DAPI and phalloidin staining images were obtained using a Nikon AXR confocal microscope and NIS Elements software (Nikon Instruments). Gel occupancy was analyzed using ImageJ software [53]. WISH images were obtained using a confocal Zeiss LSM 880, with Zen Blue 2.6 software (Zeiss Microscopy, Thornwood, NY, USA.)

---

**Supplemental Figure 1.**
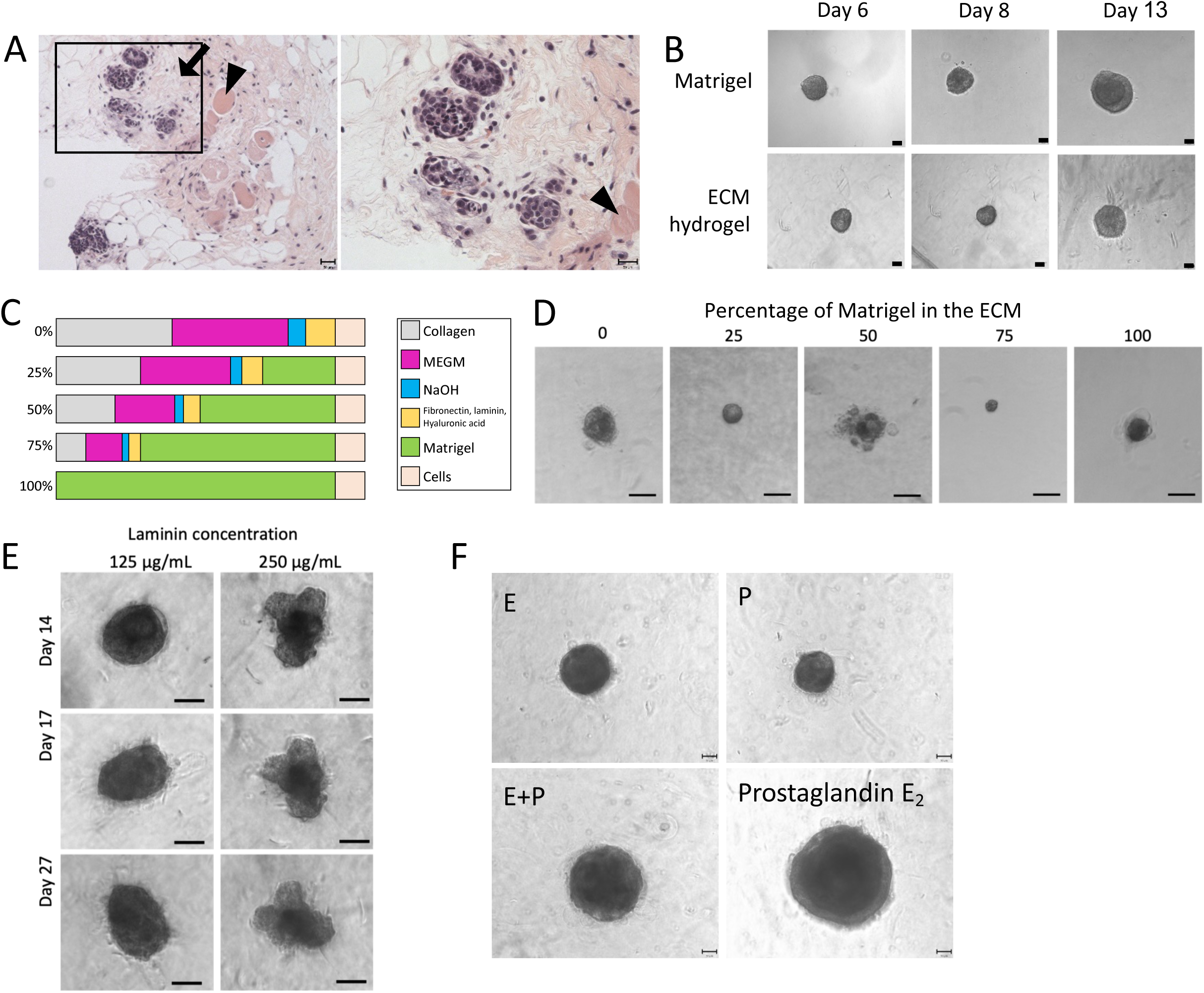
Attempts to generate opossum mammary tissue organoids. **A.** Hematoxylin-Eosin staining of an opossum mammary gland, showing the ilio marsupialis muscle (arrowheads), fibrotic tissue surrounding the epithelium (arrow). The right image is an enlargement of the area framed in the left image. **B.** Brightfield images of representative structures formed from opossum mammary epithelial cells in 3D Matrigel or ECM hydrogel over the course of 13 days. Images in sequential days are not necessarily of the same organoid. Scale bar = 50 μm. **C.** Diagram depicting the compositions of ECM hydrogel-Matrigel combination gels. **D.** Brightfield images of representative structures formed from opossum mammary epithelial cells cultured in ECM hydrogels with different percentages of Matrigel (as depicted in **B**). Scale bar = 50 μm. **E.** Brightfield images of representative structures formed from opossum mammary epithelial cells in ECM hydrogel containing two concentrations of laminin, over the course of 27 days. Images in sequential days are not necessarily of the same organoid. Scale bar = 50 μm. **F.** Brightfield images of representative structures formed from opossum mammary epithelial cells in ECM hydrogel in the presence of estrogen (“E”), progesterone (“P”), a combination of both, or prostaglandin E2. Scale bar = 50 μm.

